# Predicting clinical outcomes of SARS-CoV-2 drug treatments with a high throughput human airway on chip platform

**DOI:** 10.1101/2022.06.07.495101

**Authors:** Christine R. Fisher, Felix Mba Medie, Rebeccah J. Luu, Landys Lopez Quezada, Robert B. Gaibler, Thomas J. Mulhern, Logan D. Rubio, Elizabeth E. Marr, Elizabeth P. Gabriel, Jeffrey T. Borenstein, Ashley L. Gard

## Abstract

Despite the relatively common observation of therapeutic efficacy in discovery screens with immortalized cell lines, the vast majority of drug candidates do not reach clinical development. Candidates that do move forward often fail to demonstrate efficacy when progressed from animal models to humans. This dilemma highlights the need for new drug screening technologies that can parse drug candidates early in development with regard to predicted relevance for clinical use. PREDICT96-ALI is a high-throughput organ-on-chip platform incorporating human primary airway epithelial cells in a dynamic tissue microenvironment. Here we demonstrate the utility of PREDICT96-ALI as an antiviral screening tool for SARS-CoV-2, combining the high-throughput functionality of a 96-well plate format in a high containment laboratory with the relevant biology of primary human tissue. PREDICT96-ALI resolved differential efficacy in five antiviral compounds over a range of drug doses. Complementary viral genome quantification and immunofluorescence microscopy readouts achieved high repeatability between devices and replicate plates. Importantly, results from testing the three antiviral drugs currently available to patients (nirmatrelvir, molnupiravir, and remdesivir) tracked with clinical outcomes, demonstrating the value of this technology as a prognostic drug discovery tool.

## Introduction

As of this writing, the COVID-19 pandemic has taken over one million lives in the United States^1^,^2^, despite the availability of effective and safe vaccines^3^,^4^. A combination of new variants and waning immunity continues to take a severe toll. Therapeutic compounds capable of reducing disease severity and limiting hospitalizations and deaths are urgently needed, and yet the process for identifying treatments has been slow and arduous. Extensive clinical trials for off-label therapies have yielded few positive results^5^, and have been plagued by controversy and confusion^6^. *De novo* screens for antiviral compounds often yield hits, but the experimental compounds are rarely tested beyond *in vitro* cell culture models^7,8,9^. Currently, the only available COVID-19 antivirals include two oral medications with Emergency Use Authorizations (Paxlovid^10,11^, and Molnupiravir^12^,^13^,^14^), and one FDA-approved intravenous medication (Remdesivir^15^,^16^), with varying degrees of efficacy seen in clinical results.

A major limiting factor in the development of antiviral treatments for COVID-19 and other respiratory viral infections such as influenza is a lack of preclinical models capable of providing data predictive of human clinical responses. The most common preclinical platforms for COVID-19 include cell culture systems based on immortalized cell lines and a range of animal models (mice, hamsters, ferrets, non-human primates). Cell culture systems incorporating immortalized cell lines (e.g., Calu-3, A549, Vero E6) lack many of the key features of human lung and airway tissues (mucus formation and ciliary beating, for instance) and therefore do not recapitulate critical mechanisms and features of the tissue microenvironment. Animal models for respiratory infectious diseases^17^ are widely known to be poorly predictive of human responses^18^, and the known versus predictive susceptibilities to infection across species show wide disparities^19^. Platforms comprising human primary airway or alveolar epithelial cells in Transwells have been utilized for SARS-CoV-2 infection modeling^20,21,22^, but often have low throughput, are inconvenient to operate and incompatible with high resolution imaging, and are of limited relevance to the *in vivo* microenvironment. The abovementioned limitations have spurred intense efforts toward the development of organs-on-chips (OoC) microfluidic platforms comprising of human primary cells cultured in a dynamic and physiologically relevant microenvironment^23,24,25,26,27,28^ toward a range of disease and safety models. While the OoC field has made many exciting technological advances, and has experienced successes in various applications in drug safety and drug development, there exists a tremendous unmet need for predictive OoC airway models to fill a critical gap in preclinical therapeutic screening for COVID-19 and other emerging respiratory infections.

Key features of organ-on-chip models necessary for application to therapeutic screening for COVID-19 include: 1) fidelity of the model in recapitulating the human airway, 2) sufficient throughput to provide statistically significant data across multiple conditions in a standardized format (such as a microtiter plate), 3) compatibility with powerful, multiplexable, and complementary readouts, and 4) suitability for operation in high-containment environments. There are several reports of lung-on-a-chip models for evaluation of COVID-19 therapeutics, but they are either practically limited to pseudovirus entry^29,30^ (instead of native SARS-CoV-2 with the complete viral life cycle), or have struggled to demonstrate robust and repeatable viral replication as required to test therapeutic efficacy in a clinically-useful manner^31^. The platform^32,33^ utilized for this study, PREDICT96-ALI (Air Liquid Interface), has previously been reported toward application in modeling therapeutics against pandemic Influenza A Viruses (IAV) as well as the common cold coronaviruses hCoV-NL63 and hCoV-OC43^34^. More recently, we reported the application of the PREDICT96-ALI platform in modeling COVID-19 infections, the first successful demonstration of robust SARS-CoV-2 viral replication in a human primary cell-based OoC model^35^. That initial capability has now been expanded to an evaluation of multiple donor sources of primary cells, providing an avenue toward understanding differences in disease severity and antiviral response between different patient populations. Here we report proof-of-concept therapeutic screening studies with current treatments for COVID-19 (nirmatrelvir (a component of Paxlovid), Molnupiravir (Lagevrio), and Remdesivir (Veklury)) in an OoC model. We demonstrate antiviral efficacy in our *in vitro* model which parallels clinical experiences as measured by effects on hospitalization rates. This new capability has the potential to revolutionize the identification and evaluation of efficacious drugs for COVID-19 and other emerging pandemic infections. By introducing a more predictive human tissue-based tool into the drug screening pipeline, this technology can significantly increase the likelihood of success in preventing severe disease through timely and effective validation of new antiviral therapies.

## Results

### Antiviral efficacy in PREDICT96-ALI

To demonstrate PREDICT96-ALI’s utility as an antiviral screening tool, 5 antiviral compounds were screened at multiple doses (10, 1, and 0.1 μM) on a single PREDICT96-ALI plate. Briefly, a PREDICT96-ALI plate (**Figure 1A**) was seeded with normal human bronchial epithelial (NHBE) tissue and cultured under air-liquid interface (ALI) conditions for 3-4 weeks^34^ (**Figure 1B**). The plate was then infected with SARS-CoV-2 (**Figure 1B**) and dosed with antiviral compounds both 2 hours post infection (p.i.) and again on days 2 and 4. The antiviral effect was first determined by quantifying viral genomes from apical wash samples (**Figure 1B**) with RT-qPCR (**Figure 2**). Untreated or vehicle-treated tissues exhibited a >1000-fold increase in viral copy number between days 0 and 6, the standard viral growth kinetics for SARS-CoV-2 in PREDICT96-ALI. Significant reduction in viral titer was observed in 4 out of 5 antivirals: nirmatrelvir, molnupiravir, remdesivir, and PF-00835231 (**Figure 2A-D**), while calpeptin (**Figure 2E**) did not exhibit an antiviral effect at the concentrations tested. Nirmatrelvir was the only drug which reduced viral titer at all concentrations tested, as well as displayed significance at the earliest timepoint of 2 days post infection (dpi). PF-00835231, a precursor to Nirmatrelvir originally developed against SARS-CoV, exhibited an antiviral effect but was less efficacious than Nirmatrelvir. All data in Figure 2 was collected from a single PREDICT96-ALI plate experiment and was representative of 2 independent experiments. **Table 1** displays inter- and intra-plate variability as percent CV calculated from viral genome titers at each timepoint.

**Table 1.** Inter- and intra-plate variability from PREDICT96-ALI over experimental time course.

|  | 0 dpi | 2 dpi | 4 dpi | 6 dpi |
| --- | --- | --- | --- | --- |
| Intra-plate variability 1 | 3.1% CV | 5.0% CV | 7.1% CV | 6.0% CV |
| Intra-plate variability 2 | 2.6% CV | 4.6% CV | 12.2% CV | 11.1% CV |
| Inter-plate variability | 3.1% CV | 8.0% CV | 13.0% CV | 11.8% CV |

**Figure 1.**
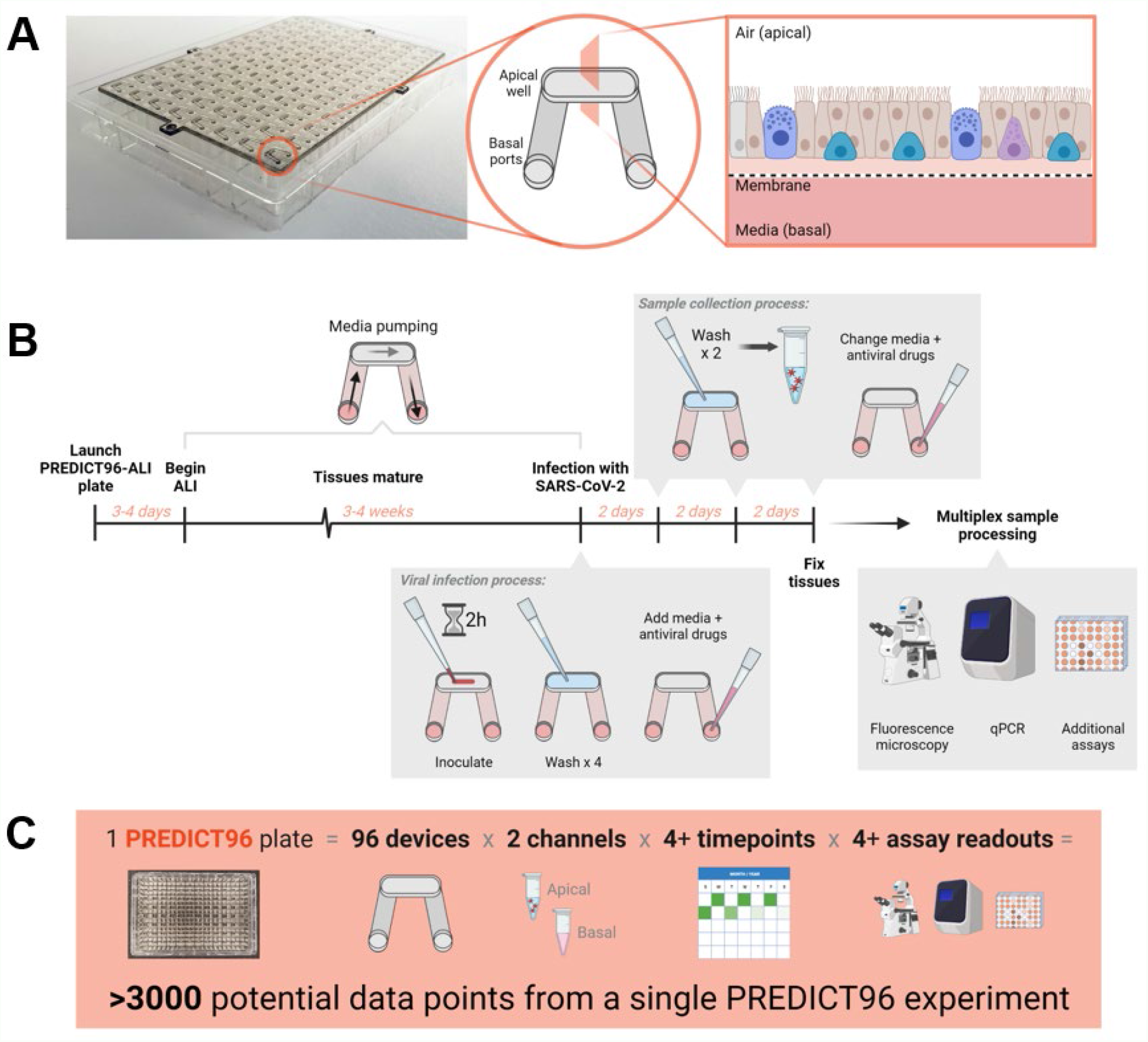
PREDICT96-ALI experimental workflow. **(A)** PREDICT96-ALI is comprised of 96 individual devices each composed of an apical well and basal channel. Lung tissue is seeded into the apical well where it is exposed to air and receives nutrients from media pumping through the basal channels. Each apical well and basal channel is separated by a porous membrane. **(B)** Lung tissue maturation in PREDICT96-ALI is detailed elsewhere^34^. After 3-4 weeks in air-liquid interface (ALI) culture, mature tissues are infected with SARS-CoV-2 for 2 hours (h), then washed with Hank’s Buffered Saline Solution (HBSS). Media containing antiviral compounds is then added to the basal channels. Every 2 days, the apical side of the tissues is washed with HBSS to collect shed virus, media is collected, and fresh media and antiviral compounds are added to the basal side. At the end of the infection, tissues are fixed with paraformaldehyde for staining and microscopy. HBSS washes are analyzed by RT-qPCR for viral genome quantification. HBSS washes and media samples can be further analyzed by additional assays. **(C)** Calculations illustrating how a single PREDICT96-ALI plate can yield >3000 data points. Figure created with BioRender.com.

**Figure 2.**
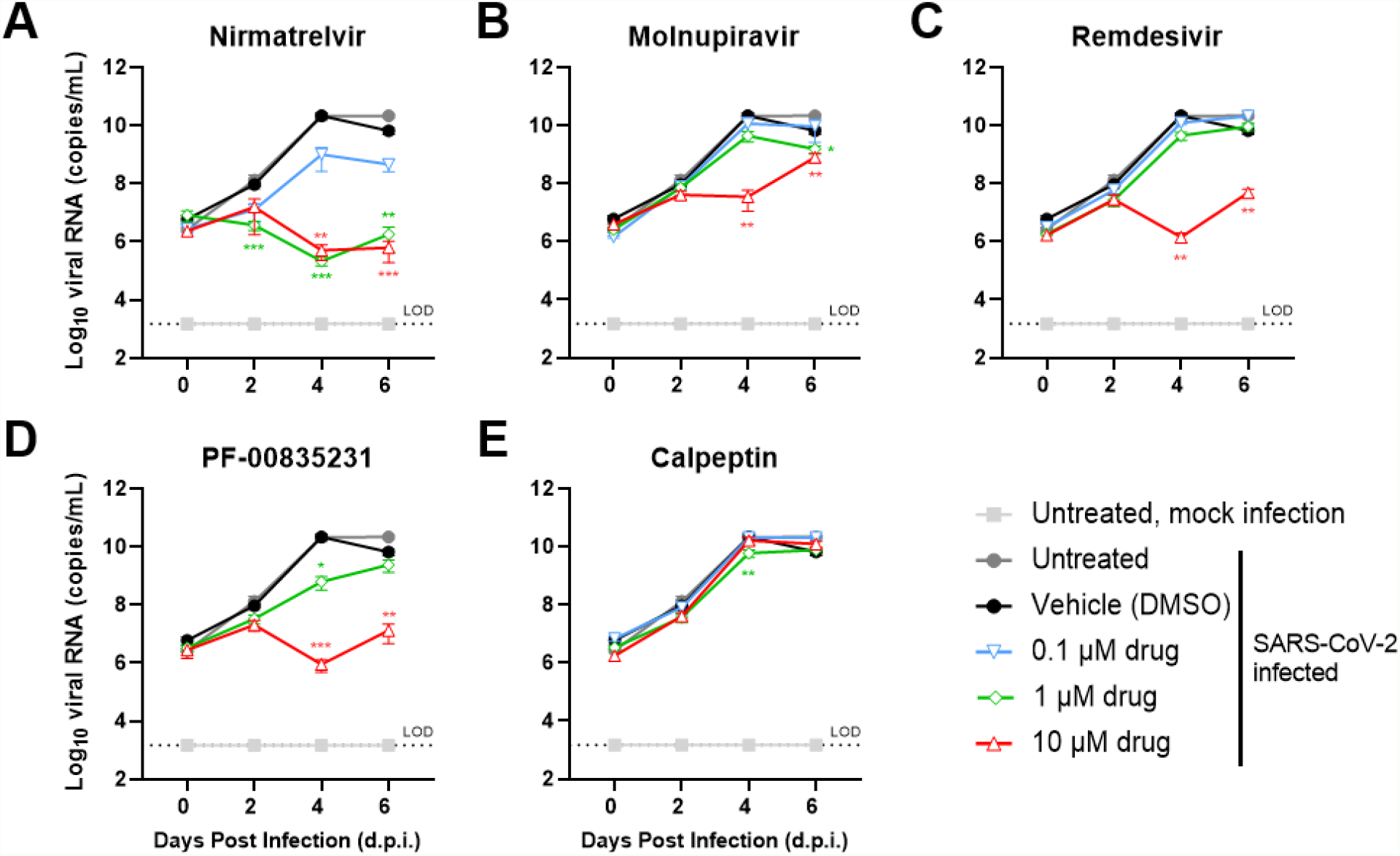
A single PREDICT96-ALI plate elucidates differential responses to antiviral drugs. Comparison between the antiviral effects of 5 drug compounds against SARS-CoV-2 infection in primary human epithelial tissue from donor 04401 grown in a PREDICT96-ALI plate: **(A)** Nirmatrelvir, **(B)** Molnupiravir, **(C)** Remdesivir, **(D)** PF-00835231, and **(E)** Calpeptin. Antiviral drugs were diluted in Pneumacult ALI medium, added 2 h after infection, and again on days 2 and 4 at the concentrations indicated. Apical sides of tissues were washed with HBSS to collect viral supernatant from which SARS-CoV-2 viral genomes were quantified by RT-qPCR, probing for the N1 gene target. N=4 tissue devices per condition (except untreated samples, n=8). Data collected from the same experiment displayed in **Figure 3** and representative of 2 independent experiments. Statistical significance determined by a two-way analysis of variance (ANOVA) with Dunnett’s test for multiple comparisons (comparing to infected/untreated): P > 0.05; *, P ≤ 0.05; **, P ≤ 0.01; ***, P ≤ 0.001; ****, P ≤ 0.0001. Unlabeled lines indicate no significant different from infected/untreated (gray) line.

### Fluorescent image quantification of antiviral screen

To complement the viral genome quantification data, the lung tissues were also fixed at 6 dpi, stained for SARS-CoV-2 nucleocapsid (N), and imaged using the Celigo Imaging Cytometer (**Figure 3**). Fluorescence data was quantified as nucleocapsid positive cells. The trends in viral titer reduction from antiviral effect match the corresponding 6 dpi RT-qPCR data (**Figure 2**). For example, Niramtrelvir was the best-performing antiviral with total reduction in nucleocapsid staining at both 10 and 1 μM (**Figure 3A**), and Calpeptin was the least-efficacious compound tested (**Figure 3E**). All data in **Figure 3** was collected from a single PREDICT96-ALI plate experiment and was representative of 2 independent experiments. These data highlight high-throughput fluorescent imaging as a complementary, protein-based readout to the nucleic acid-based RT-qPCR assay and demonstrate the robustness of the platform in its capability to screen antiviral compounds.

**Figure 3.**
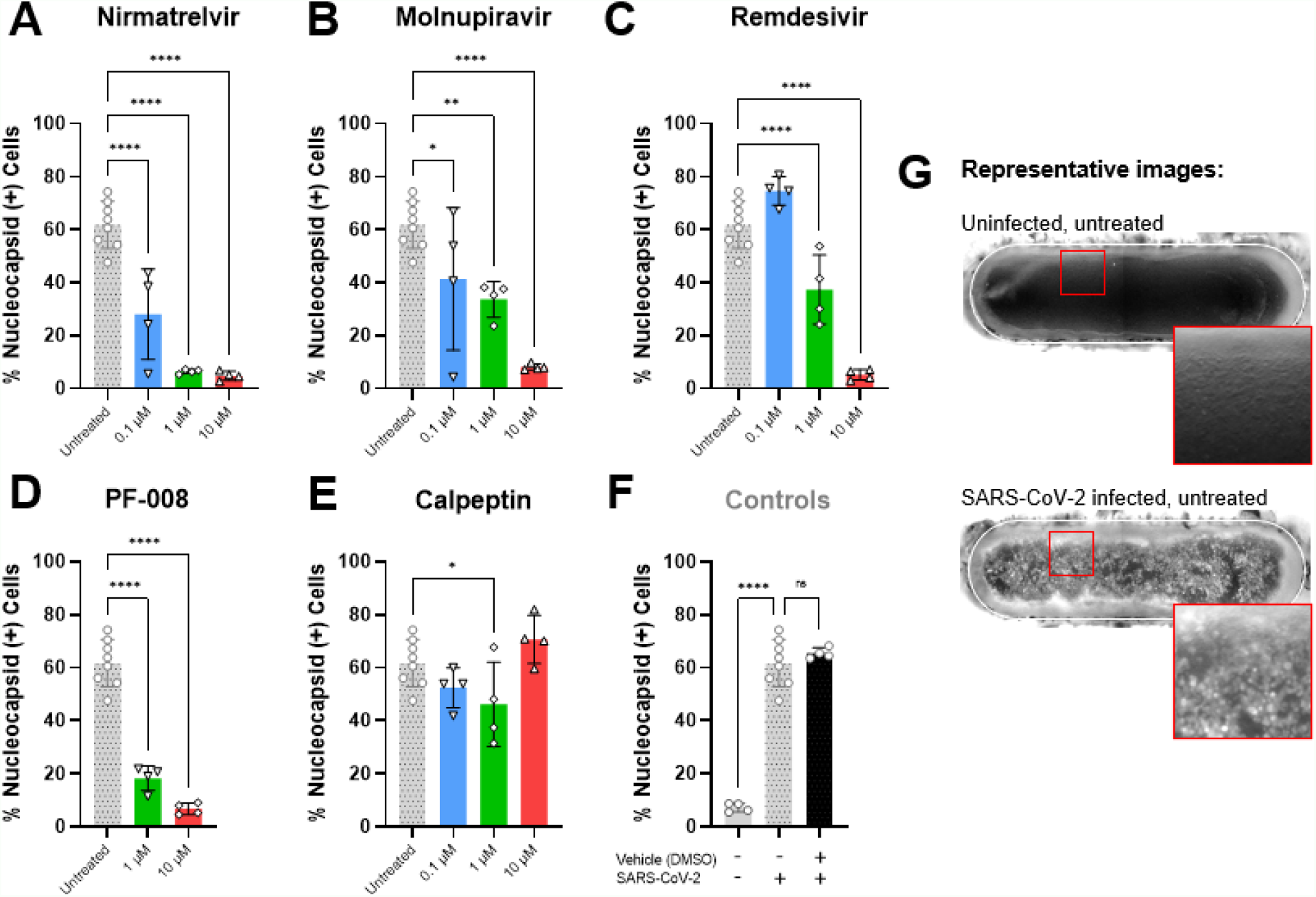
High-throughput fluorescence imaging data complements antiviral drug screen. Image quantification of SARS-CoV-2 infected tissues nucleocapsid positive (+) cells shows tissue response to 5 drug compounds 6 days post inoculation of SARS-CoV-2 infection in primary human epithelial tissue from donor 04401 grown in a PREDICT96-ALI plate: **(A)** Nirmatrelvir, **(B)** Molnupiravir, **(C)** Remdesivir, **(D)** PF-00835231, and **(E)** Calpeptin. Briefly, tissues were stained for nuclei (DAPI) and SARS-CoV-2 nucleocapsid (N) protein. Using the Celigo software, the percentage of nuclei corresponding to cells positive for the nucleocapsid protein was quantified. **(F)** Controls: Untreated, uninfected (solid gray bar), untreated, infected (patterned bar), and vehicle treated, infected (black bar). **(G)** Representative monochromatic fluorescence images of uninfected and SARS-CoV-2-infected tissues stained for SARS-CoV-2 nucleocapsid. All data in this figure collected from the same experiment displayed in **Figure 2** and representative of 2 independent experiments. Bar graphs show mean ± standard deviation; n=4 tissue devices per condition (except untreated samples, n=8). Statistical significance determined by a one-way analysis of variance (ANOVA) with Dunnett’s test for multiple comparisons (comparing to infected/untreated): no significance (ns), P > 0.05; *, P ≤ 0.05; **, P ≤ 0.01; ***, P ≤ 0.001; ****, P ≤ 0.0001. Unlabeled bars indicate no significance.

### PREDICT96-ALI supports multiple tissue donors

To further demonstrate the powerful capability that a high-throughput primary tissue model confers, PREDICT96-ALI was seeded with tissues from 4 healthy human donors and infected with SARS-CoV-2 following the same infection scheme in **Figure 1B**. All four donor tissues were permissible to infection with SARS-CoV-2 and replicated the virus to high titers (**Figure 4A**). From day 0 to day 6 p.i., viral titers increased 1502-fold, 2914-fold, 8579-fold, and 966-fold by donors 05101, 04282, 04401, 06288, respectively. Representative immunofluorescence images for each donor show robust infection of ciliated cells within the pseudostratified lung tissue via SARS-CoV- 2 nucleocapsid staining (**Figure 4B**). The ability to test tissues from diverse human patient populations is a key feature of high throughput organ-on-chip technologies. In future studies, PREDICT96-ALI’s throughput and reproducibility can be applied to antiviral drug screening on diverse patient populations to yield insights on individual responses to candidate drugs.

**Figure 4.**
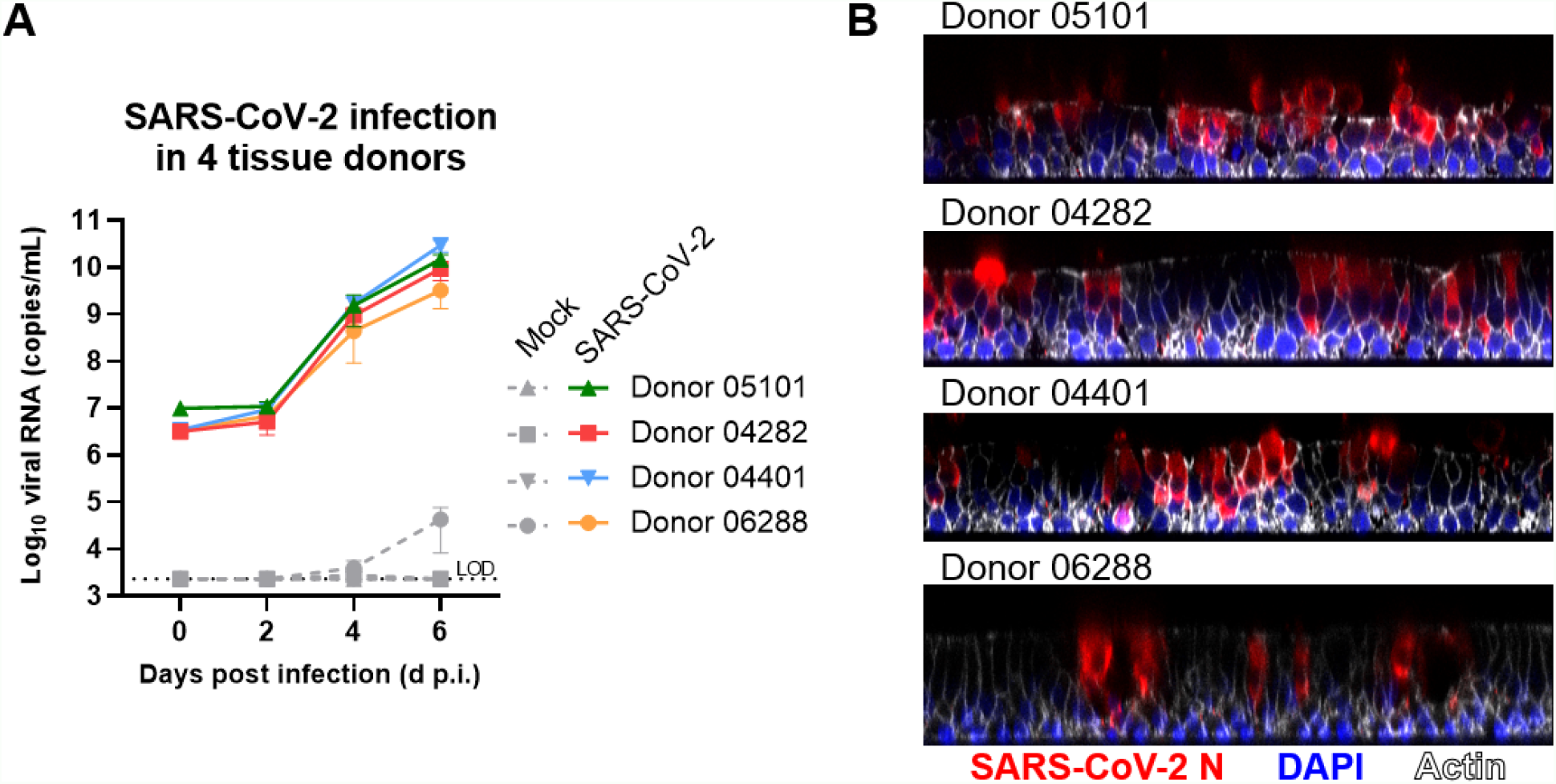
PREDICT96 supports SARS-CoV-2 infection in multiple tissue donors. Infection kinetics of SARS-CoV-2-inoculated PREDICT96-ALI airway tissues. **(A)** PREDICT96-ALI airway tissues generated from four human donors and inoculated with SARS-CoV-2 USA-WA1 showed an increase in viral load via RT-qPCR between 2 and 6 days post infection (d p.i.). The following demographic information pertains to the four donors tested: donor 05101, 29 year old Caucasian female (non-smoker); donor 04282, 23 year old Caucasian male (marijuana smoker); donor 04401, 16 year old Caucasian male; donor 06288, 70 year old Caucasian female (non-smoker). Mean ± standard error of the mean shown (n=3 for mock samples, n=6 for SARS-CoV-2-infected samples). Statistical significance determined by a two-way analysis of variance (ANOVA) with Tukey’s test for multiple comparisons: no significance detected between infected donors at 2, 4, or 6 d p.i. **(B)** Orthogonal immunofluorescence images of pseudostratified lung tissues generated from four NHBE donors. Tissues were stained with an antibody for SARS-CoV-2 nucleocapsid (N, red), nuclei (DAPI, blue), and phalloidin (actin, white).

## Discussion

More than two years into the COVID-19 pandemic, antiviral therapeutics against SARS-CoV-2 remain limited in number, availability and effectiveness. Progress towards development and approval of SARS-CoV-2 drugs has been hindered by inaccurate guidance stemming from efficacy screens in oversimplified cell culture systems^29^. Consequently, precious resources in animal and human clinical trials are spent on drug candidates which have rarely been vetted on primary human tissue. Data gained from using proxies for human tissue (e.g. immortalized cell lines) or SARS-CoV-2 (e.g. pseudoviruses) do not reliably yield predictive data making drug candidates more likely to fail in clinical trials^36^. Furthermore, such systems are rarely able to address a drug candidate’s efficacy in varied patient populations.

Draper has built the first model of the human lung capable of assessing efficacy of SARS-CoV-2 therapeutics in high-throughput. The PREDICT96-ALI system accurately recapitulated the wins and fails of the three FDA EUA-approved antiviral therapeutics: both viral genome and nucleocapsid staining data (**Figures 2 and 3**) tracked with clinical outcomes documented by reduction in hospitalization time (**Table 2**).

**Table 2.**
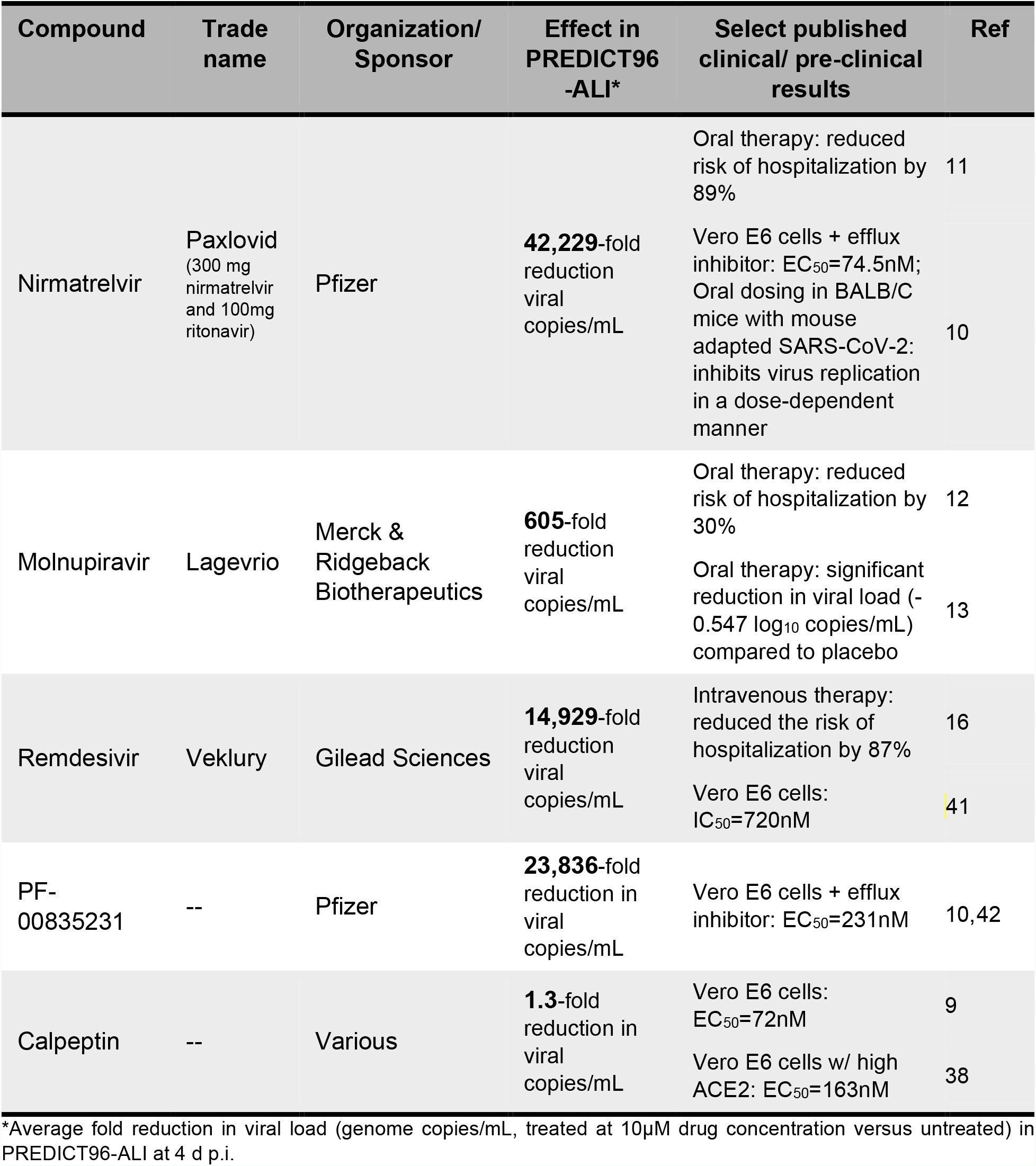
Comparison of Published Clinical Outcomes and Viral Load Reduction in PREDICT96-ALI.

The compounds tested here are from three classes of drugs with distinctive mechanisms: Nirmatrelvir and PF-00835231 are SARS-CoV-2 main protease (M^pro^) inhibitors which bind to the active site of M^pro^ and prevent viral entry^10^; Molnupiravir is a ribonucleoside analog of cytidine^14^ and Remdesivir is a ribonucleotide analogue of adenosine monophosphate (AMP)^37^, both of which introduce errors into the RNA genome during SARS-CoV-2 replication. Calpeptin is a calpain inhibitor which has also been shown to bind to the active site of M^pro 9^. PREDICT96-ALI was able to detect antiviral efficacy across drug classes, and also resolved significant differences between members of the same class.

Calpeptin, the least efficacious compound tested in this study, has been identified in several SARS-CoV-2 drug screens^9, 38,39^, even showing the highest antiviral activity in a screen based on x-ray crystallography and a SARS-CoV-2 screen on Vero E6 cells^9^. Despite this, no studies have reported its efficacy in animal models or clinical trials and its lack of efficacy in PREDICT96-ALI likely predicts its failure as a SARS-CoV-2 therapeutic. Conversely, the best-performing drug tested in PREDICT96-ALI, Nirmatrelvir, was tested in an NHBE model in addition to the cell lines Vero E6 and A549 during its preclinical development^10^. The EC90 from the NHBE assay was furthermore selected as the basis for setting the minimum systemic unbound plasma concentration to be maintained in animal models and clinical trials. Given the clinical success of Paxlovid (Nirmatrelvir/Ritonavir), the use of a lung model to not only test but base clinical studies on serves as a powerful validation for the utility of human microphysiological systems.

PREDICT96-ALI represents a powerful advancement in its high-throughput capacity which enables statistical significance while maintaining robust biological function. The 96-well form factor (**Figure 1A**) and low inter-plate variability (**Table 1**) allow for a balance between minimizing numbers of biological replicates and maximizing numbers of conditions in a given experiment. Furthermore, low intra-plate variability (**Table 1**) enables comparison between plates in different experiments, a key factor PREDICT96-ALI’s scalability.

Here we have presented two complementary assay readouts, RT-qPCR on apical wash samples and fluorescence staining on fixed tissues, which were performed using materials from the same PREDICT96-ALI plate. As RT-qPCR to detect viral genomes uses only 4% of the apical wash sample (8μl out of 200 μl collected total), there is ample remaining volume of apical wash samples to perform additional assays such as live virus titering or immunoassays to assess disease state or host response. Additionally, basal media samples accumulate valuable analytes which can be assayed in a physiologically relevant concentration. Altogether, PREDICT96-ALI can potentially yield >3000 unique datapoints (**Figure 1C**), turning a single experiment into a trove of data to characterize a candidate therapeutic’s effect. In the context of antiviral therapeutics, data from such complementary assays can contribute to mechanistic studies by capturing the state of infection at several points along the viral life cycle where inhibition can occur. Ultimately, PREDICT96-ALI can significantly impact disease prevention through the timely and effective screening of new antiviral therapies.

## Methods and Materials

### NHBE culture

Primary normal human bronchial/tracheal epithelial cells (NHBE) (Lifeline Cell Technology) were expanded in BronchiaLife Epithelial Airway Medium (Lifeline Cell Technology) and passaged once prior to cryopreservation in Cryostor CS10 (STEMCELL). NHBEs were seeded in PREDICT96- ALI plates as previously described^34^. The following demographic information pertains to the four donors tested: donor 05101, 29 year old Caucasian female (non-smoker); donor 04282, 23 year old Caucasian male (marijuana smoker); donor 04401, 16 year old Caucasian male; donor 06288, 70 year old Caucasian female (non-smoker). NHBEs were thawed and seeded immediately at 10,000 cells per device in Small Airway Growth Medium (CC-3118, Lonza) supplemented with 5 μM Y-27632 (Tocris), 1 μM A 83-01 (Tocris), 0.2 μM DMH-1 (Tocris), 0.5 μM CHIR 99021 (Tocris), and 1x penicillin-streptomycin (P4333, Sigma). After 24-48 h of culture, media was replaced with Air-Liquid Interface Epithelial Differentiation Medium (Lifeline Cell Technology). After 48 h of culture, tissues were lifted into air-liquid interface (ALI) culture by aspirating all media from the apical well of each device and replacing the media in the basal channel with Pneumacult-ALI (STEMCELL). Flow was initiated at 1 μL/m in the basal channels via the PREDICT96 pump^32^. Tissues were washed once per week with Hanks Balanced Salt Solution (HBSS, Sigma). Tissues were cultured in ALI at 37°C in 5% CO_2_ and 90% humidity for 3-4 weeks prior to infection. Media in the basal channels was refreshed 6 days a week.

### Infection with SARS-CoV-2

All SARS-CoV-2 infections were performed in a Biosafety Level 3 (BSL-3) facility at the New England Regional Biosafety Laboratory at the Tufts University with appropriate institutional biosafety committee approvals. SARS-CoV-2 isolate USA-WA1/2020 passage 4 virus was obtained from Joseph Brewoo and Sam R. Telford III (New England Regional Biosafety Laboratory, Global Heath & Infectious Disease, Tufts University). A passage 5 stock for USA- WA1/2020 strain was generated by propagating the virus on Vero E6 cells (ATCC) in high-glucose Dulbecco’s Modified Eagle Medium (DMEM, Gibco) supplemented with 0.1% bovine serum albumin (Sigma), 1x Penicillin/Streptomycin (Gibco), 1x non-essential amino acids (NEAA, Gibco). Inoculation of PREDICT96-ALI airway tissues with SARS-CoV-2 (and mock control) were performed similarly to coronavirus infections described previously^34^. Briefly, the PREDICT96-ALI tissues were prepared by first washing tissue with Hanks’ Balanced Salt Solution (HBSS, Sigma) to remove mucous. Freshly-thawed SARS-CoV-2 was diluted in the same infection media used for viral propagation: 8 × 10^4^ pfu (**Figure 3**) or 1 × 10^5^ pfu (**Figure 4**) total viral input. Mock infections were performed by adding the same volume of infection media without virus to the apical side of devices. Viral inoculum (or mock control) was incubated on the apical surface of the microtissues for 2 h rocking at 34°C. Inoculum was subsequently removed, and the apical chambers of each microtissue device were washed 4x with HBSS. The final wash was reserved as the sample representing 0 days post infection (d p.i.). Fresh Pneumacult-ALI media (with or without therapeutics, further detailed in Antiviral Drug Dosing section) was introduced into the basal channels before static incubation at 34°C. On 2, 4, and 6 d p.i., the apical surface of the tissues was washed twice with HBSS (40 m per wash). The two HBSS washes collected from the apical surface of a single device were pooled and stored at -80°C until downstream assessment. Fresh Pneumacult-ALI media was introduced into the bottom chamber before returning the plate to 34°C. At the conclusion of the experiment on day 6 p.i., the PREDICT96-ALI airway tissues were fixed with 4% paraformaldehyde (Electron Microscopy Sciences) for 1 h at room temperature. Tissues were then washed with phosphate buffered saline (PBS) and stored at 4°C (submerged in PBS) until immunofluorescence staining.

### Antiviral drug dosing

Nirmatrelvir (MedChemExpress), PF-00835231 (MedChemExpress), Molnupiravir (Tocris Bioscience), Calpeptin (Tocris Bioscience) and Remdesivir (Tocris Bioscience) were used in anti- viral screens performed on PREDICT96-ALI tissue. All compounds were diluted fresh from 10 mM stock concentrations in dimethyl sulfoxide (DMSO) to 10, 1 or 0.1 µM in complete Pneumacult-ALI. DMSO was diluted similarly to 10 μM to serve as a vehicle control. Drugs or DMSO diluted in Pneumacult-ALI were applied to select bottom channels of PREDICT-ALI airway tissues 2 h after infection and on days 2 and 4 p.i. All drug dosing conditions were performed in quadruplicate, accounting for 60 devices of the PREDICT96-ALI plate; 16 additional devices accounted for biological controls; 8 devices accounted for assay control. The plate can accommodate testing 1 additional compound in this experimental configuration.

### RT-qPCR

HBSS washes from PREDICT96-ALI devices were tested by reverse transcription quantitative polymerase chain reaction (RT-qPCR)^40^. Briefly, aliquots of wash samples from PREDICT96-ALI devices were inactivated at 70°C for 10 m in a MiniAmp Thermal Cycler (Applied Biosystems) and stored at -80°C until use. Direct RT-qPCR was then performed on heat-inactivated samples using the QIAprep&amp Viral RNA UM Kit (Qiagen) with 8 μl heat inactivated sample in a 20 μl reaction. The reaction was run in an Applied Biosystems QuantStudio 7 Flex System (Thermo Scientific) using the following condition: 50°C for 10 m, 95°C for 2 m, 40 cycles of 95°C for 5 s and 60°C for 30 s. The following primers and probes targeting SARS-CoV-2 nucleocapsid protein were used to detect viral RNA in HBSS wash samples: nCOV_N1 Forward Primer Aliquot, 100 nmol (IDT), nCOV_N1 Reverse Primer Aliquot, 100 nmol (IDT), nCOV_N1 Probe Aliquot, 50 nmol (IDT). Samples with cycle threshold (ct) values above 37 were omitted. Absolute quantification (copies/mL) of washes viral RNA was calculated using a standard curve generated from serial dilutions of 2019-nCoV_N_Positive Control viral RNA (IDT).

### Immunofluorescence staining

Tissues were permeabilized with 0.3% Triton X-100 (Sigma) for 30 m and then incubated in blocking buffer consisting of 3% normal goat serum (Thermo Fisher) and 0.1% Tween-20 (Sigma) in PBS for 1 h at room temperature, rocking. SARS-CoV-2 nucleocapsid (N) primary antibody (Genetex) was diluted 1:100 in blocking buffer and tissues incubated overnight at 4°C, rocking. Tissues were washed twice with 0.1% Tween-20 and then once with blocking buffer. Alexa Fluor Secondary antibody Alexa Fluor 555 goat anti-rabbit (Thermo Fisher) was diluted 1:300 in blocking buffer and incubated overnight at 4°C, rocking. Tissues were washed three times with 0.1% Tween-20. Hoechst 33342 (Thermo Fisher) Phalloidin-iFluor 647 (Abcam) were diluted 1:800 and 1:1000, respectively, in 0.1% Tween-20 and incubated at room temperature for 30 m, rocking. Tissues were rinsed with PBS three times and then stored in PBS until imaging.

### Fluorescence imaging

For high-throughput fluorescence imaging, 10x magnified images of tissues fixed at 6 d p.i. were captured using a custom 96 well plate map created for the PREDICT96-ALI dimensions on the Celigo Imaging Cytometer (Nexcelom). Using the Celigo software (version 5.3), the nuclei from each device were located from the blue channel and outlined as a region of interest (ROI). Nuclei at the edges of the device were gated out to account for edge effects. To quantify the percentage of nucleocapsid (N)-positive cells, co-localization of N expression of each ROI was determined. If the signal intensity of N in the red channel was greater than the background signal of uninfected controls, the ROI was determined as positive. To calculate percent positivity, the number of nucleocapsid positive cells was divided by the total number of nuclei in the device. Results from each condition were averaged.

For confocal imaging, stained tissues were imaged using a Zeiss LSM700 laser scanning confocal microscope and Zen 2012 SP5 (black edition) software. Z-stacks of the tissues were acquired with a 40x objective.

### Statistical analysis

Statistical analysis was performed in Prism software version 9.3.0 (Graphpad). RT-qPCR data (Figures 2 and 3) were log-transformed and analyzed using a two-way analysis of variance (ANOVA) with either Dunnett’s or Tukey’s test for multiple comparisons (noted in figure legend). Mean fluorescence intensity (MFI) as a measure of percent nucleocapsid positive cells (Figure 4) was analyzed using a one-way ANOVA with Dunnett’s test for multiple comparisons. All P-values signified as follows: P > 0.05, no significance (ns); P ≤ 0.05, *; P ≤ 0.01, **; P ≤ 0.001, ***; P ≤ 0.0001, ****. For both RT-qPCR and MFI data, the Shapiro-Wilk test did not show evidence of non-normality.

## Acknowledgements

The authors gratefully acknowledge Roger Odegard, Timothy Petrie, Robert Larsen, Jonathan Cash, David O’Dowd, Rachel Fezzie, and Alla Gimbel, for their technical and programmatic support for this work, Richard Crocker and Karen Arenburg for support of the laboratory infrastructure, John Gilmartin from the Draper Safety Office, Hesham Azizgolshani, Brian Cain and Yazmin Obi for technical support, and Corin Williams and Else Vedula for key technical guidance and critical review of the manuscript. We gratefully acknowledge the technical and programmatic support of Sam Telford, Joseph Brewoo and the staff at the Tufts University New England Regional Biosafety Laboratory. The following reagent was deposited by the Centers for Disease Control and Prevention and obtained through BEI Resources, NIAID, NIH (via Sam Telford and Joseph Brewoo of Tufts University): SARS-Related Coronavirus 2, Isolate USA- WA1/2020, NR-52281.

